# Atypical memory B cells contribute to heterogeneous vaccine responses in ocrelizumab-treated patients with multiple sclerosis

**DOI:** 10.1101/2025.11.03.686372

**Authors:** Ryan Curtin, Yogambigai Velmurugu, Fatoumatta Dibba, Yuan Hao, Chaitra Sreenivasaiah, Alireza Khodadadi-Jamayran, Samantha Nyovanie, Angie Kim, Marie L Samanovic, Mark Mulligan, Jessica Priest, Mark Cabatingan, Ryan C. Winger, Yury Patskovsky, Ilya Kister, Gregg J. Silverman, Michelle Krogsgaard

## Abstract

**Background and Objectives:** Patients with multiple sclerosis (pwMS) treated with ocrelizumab (OCR), a B-cell-depleting therapy, exhibit heterogeneous humoral responses to SARS-CoV-2 mRNA vaccination. The mechanisms underlying this variability remain incompletely understood. We performed a longitudinal analysis of B-cell subset dynamics and antigen-specific T cell responses in OCR-treated pwMS and healthy controls to determine how immune cell composition, timing of OCR infusion, and lymphocyte dynamics influence humoral response outcomes.

**Methods:** Based on post-vaccination anti-Spike IgG titers measured by multiplex bead immunoassay, pwMS were categorized as super-responders (SR), responders (R), or non-responders (NR). A 35-marker spectral flow cytometry panel was used to characterize T- and B-cell subsets longitudinally, at baseline and following stimulation with a SARS-CoV-2 peptide pool.

**Results:** CD4^+^ and CD8^+^ T cell populations were preserved across OCR-treated pwMS, and SARS-CoV-2-specific T cells remained detectable for more than 6 months after vaccination. In contrast, residual B-cell subset composition distinguished responders from non-responders. DN2-like B cells (CD19^+^CD27-IgD-T-bet^+^CD11c^+^CXCR5-) persisted despite repeated OCR infusions and were enriched in SR compared with NR. Repletion of mature naïve B cells in peripheral blood correlated with time since last OCR infusion and with stronger humoral immune responses.

**Conclusions:** B-cell subsets that resist OCR-mediated depletion may contribute to vaccine responsiveness in OCR-treated pwMS. Altered repletion kinetics of naïve B cell subsets in non-responders suggest that specific B-cell populations may serve as predictive biomarkers of vaccine-induced humoral immunity, despite preserved T-cell responses.

## Introduction

Individuals receiving immunosuppressive disease-modifying therapies (DMTs) for autoimmune conditions such as multiple sclerosis (MS) often exhibit impaired humoral and/or cellular responses to SARS-CoV-2 mRNA vaccination (1–8). Ocrelizumab (OCR), a humanized, monoclonal anti-CD20 (aCD20) antibody, is a widely used DMT for MS. OCR reduces relapse rates, limits MRI-detected lesion formation, and slows disease progression, likely through depletion of autoreactive CD20^+^ B cells and a subset of CD20^+^ T cells involved in central nervous system (CNS) inflammation (9–11). Many studies have shown that cellular immune responses to SARS-CoV-2 vaccination are preserved in OCR-treated pwMS. However, B-cell depletion is associated with markedly reduced antibody responses to mRNA vaccines, with substantial interindividual variability that remains unexplained (8, 12, 13).

In healthy individuals, transitional B cells generated in the bone marrow enter circulation and differentiate into mature naïve B cells, which can further develop into antibody-producing cells following infection or vaccination (14–16). In OCR-treated pwMS, however, transitional, mature naïve, and most CD20^+^ memory B cells are depleted from the peripheral circulation, raising the question of how some patients mount vaccine-induced humoral immune responses (9). One possibility is that CD20^+^ B cells in secondary lymphoid organs escape depletion and contribute to subsequent vaccine responses (17, 18). In addition, long-lived plasma cells lack CD20 expression and reside in protective bone marrow niches, allowing for continued antibody production despite OCR-therapy (14, 19). Prior studies also suggest that recirculating IgA^+^ plasmablasts may be resistant to depletion by anti-CD20 therapies and persist following OCR infusion (18–21).

Emerging evidence indicates that a subset of atypical CD19^+^ memory B cells, termed double-negative (DN) B cells because they lack CD27 and IgD expression, contribute to adaptive immune responses (22, 23). Among these, DN2-like B cells are characterized by CD11c surface expression and intracellular expression of the transcription factor T-bet and develop via an alternative extrafollicular pathway independent of germinal centers, enabling rapid production of hypermutated antibodies (22–25). Studies in rheumatoid arthritis and systemic lupus erythematosus demonstrate that DN2 cells share B-cell receptor (BCR) clonotypes with antibody-secreting cells (ASC), supporting their role as precursors (23, 25–27). However, their susceptibility to OCR-mediated depletion remains unclear, with conflicting reports regarding CD20 surface expression (23, 27, 28). Despite growing interest in this cell population, the role of DN2 B cells in vaccine-induced humoral immune responses in the context of B cell-depleting therapy remains incompletely understood.

Here, we performed a longitudinal analysis of B- and T-cell subsets in OCR-treated pwMS, categorized as super-responders (SR), responders (R), and non-responders (NR) based on anti-spike IgG titers following SARS-CoV-2 vaccination. Using spectral flow cytometry, dimensionality reduction, and unsupervised hierarchical clustering, we assessed antigen-specific T cell phenotypes and their relationships with heterogeneity in humoral responses up to 6 months after vaccination. We also quantified the time since the last OCR infusion for each sample, to characterize B-cell subset reconstitution across treatment cycles and evaluate how these dynamics relate to variability in vaccine-induced humoral immunity.

## Results

### Participant enrollment

To investigate the mechanisms influencing immune responses in pwMS treated with OCR, we analyzed a total of 66 patients from two combined studies, with sample collection occurring at baseline (pre-vaccination), and 4 -24 weeks after SARS-CoV-2 vaccination (Figure 1A) (12). All participants were recruited from the NYU MS Care Center, ensuring standardized infusion and sample-collection protocols. We assessed humoral and cellular immune responses to SARS-CoV-2 vaccination at predefined time points following vaccine doses 1, 2, and booster administration (Figure 1A) (8). Sample collection dates were mapped relative to the most recent OCR infusion to evaluate effects on circulating B and T cell subsets (Figure 1A). Patient demographics are shown in Supplementary Table 1. Healthy control samples were obtained from the NYU Vaccine Center.

**Figure 1:**
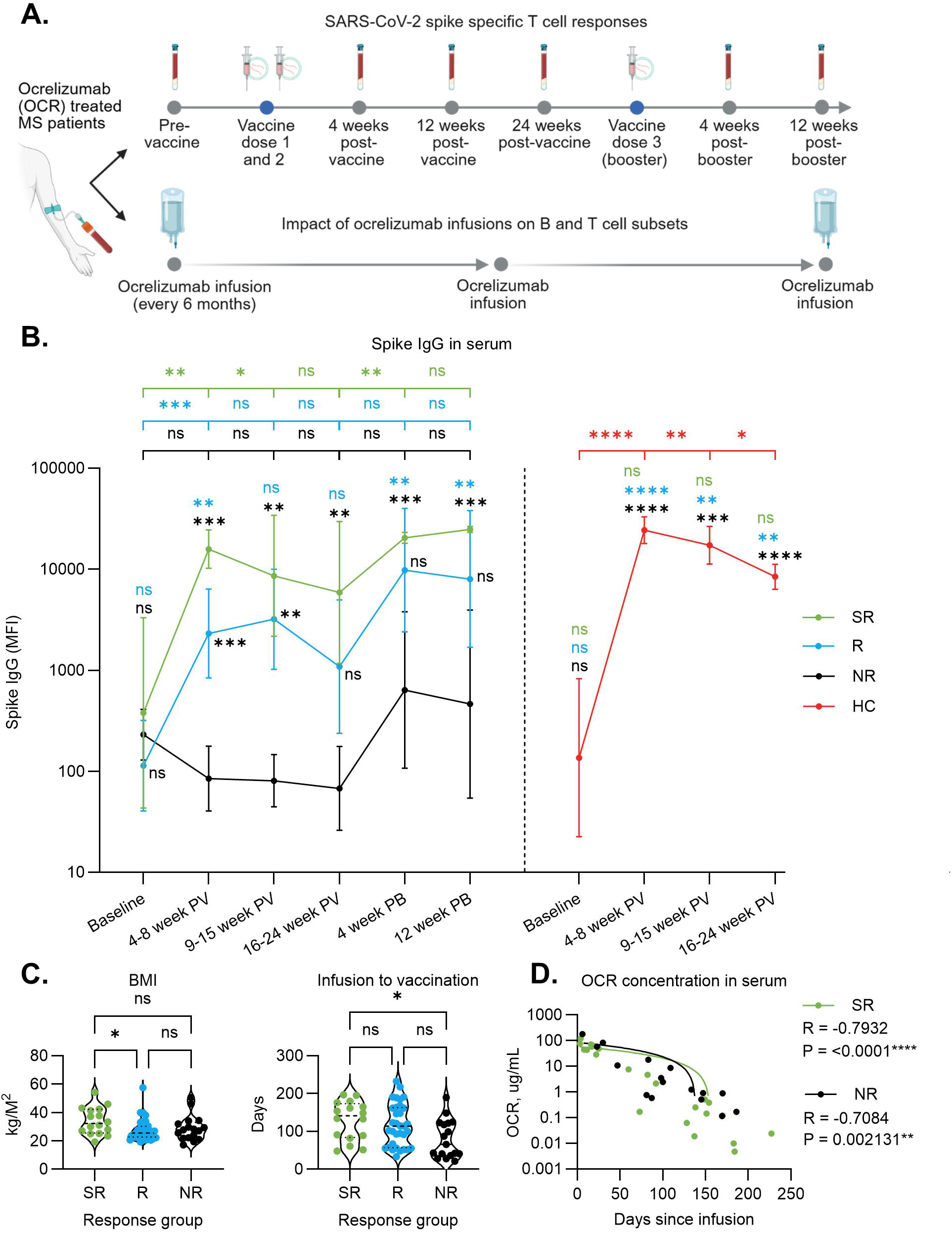
Ocrelizumab-treated pwMS show heterogeneous Spike IgG production following SARS-CoV-2 mRNA vaccination. **(A)** Study design and sample collection of OCR-treated pwMS enrolled in the longitudinal VIOLA study (n=30, top timeline). Time since most recent OCR infusion was calculated based on each patient’s post-vaccination collection dates relative to their pre- and post-vaccination infusions, which occurred approximately every six months (bottom timeline). Additional OCR-treated pwMS from a cross-sectional study (n=36) were included for post-infusion timeline analysis. Details of both studies can be found under “patients and samples”. **(B)** Serum anti**-**Spike IgG median fluorescence intensity (MFI) responses in OCR-treated pwMS and healthy controls. OCR-treated pwMS are further divided into SR (green, n=16), R (blue, n=34), and NR (black, n=16), based on quartile analysis of the earliest post-vaccination time point for each patient. Healthy controls are shown separately in red (n=10) **(C)** Comparison of OCR-treated pwMS (SR: green, R: blue, NR: black) body-mass index (BMI) at baseline (left), and days from infusion to vaccination (right). **(D)** Serum OCR concentrations compared between a subset of OCR-treated pwMS in the SR (green) and NR (black) groups. In (B) only, a mixed-effects model with the Geisser-Greenhouse correction followed by a post-hoc Tukey’s Honest Significant Difference (HSD) test was used to correct for false discovery rate (FDR). In (B), Statistics located above the graph reflect longitudinal intra-group comparisons, while statistics on the graph itself (directly adjacent to the data points) reflect inter-group comparisons at each timepoint. The color of each statistical comparison corresponds to the group to which it is compared. Additional statistical testing was performed using the Kruskal-Wallis test followed by Dunn’s post-hoc test for multi-group comparisons or Spearman’s correlation for analyses of OCR concentration. p<0.05 (*), p<0.01 (**), p<0.001 (***), and p<0.0001 (****). NS= not significant.

### Heterogeneity of humoral responses to the SARS-CoV-2 vaccination in OCR-treated pwMS

Patients were stratified into groups based on post-vaccination anti-spike IgG levels (measured as median fluorescence intensity (MFI)), using the earliest available time point for each patient (Figure 1B). Quartiles, defined as follows, were used to assign patients to groups: Q1 (MFI < 250) ‘non-responders’ (NR); Q2-Q3 (MFI 251-8884) ‘responders’ (R); and Q4 (MFI > 8885) ‘super-responders’ (SR) (Figure 1B). All healthy controls were classified as SR (Figure 1B).

To analyze the impact of demographic variables on humoral responses to vaccination, we compared patient demographics across the three response groups. The SR group had a higher body mass index (BMI) than the R group (Figure 1C). Time from OCR infusion to SARS-CoV-2 vaccination was shorter in the NR group compared with SR (Figure 1C). However, this difference alone is unlikely to fully account for the observed heterogeneity in humoral immune responses, given that the percentage of peripheral CD19^+^ B cells at each patient’s baseline collection was similar across all groups (Supplementary Figure 1A). Serum OCR concentrations declined over time with similar negative correlations between time since infusion and OCR levels in SR and NR groups (Figure 1D). Other variables were comparable across the three subgroups (Supplementary Table 1, Supplementary Figure 1B). The overall similarity in clinical demographics between the response groups suggests that an immunological mechanism may underlie the observed heterogeneity in humoral responses to vaccination.

### Antigen-specific CD4^+^ T cell profiles are comparable between pwMS and healthy controls

We evaluated the frequency of SARS-CoV-2 Spike-specific CD4^+^ T cells across OCR-treated pwMS groups and healthy controls. Antigen-induced marker-positive (AIM^+^) CD4^+^ T cells were defined by co-expression of CD137 and OX40 following *ex vivo* stimulation with SARS-CoV-2 spike peptide pools.

The complete set of phenotyping markers used in the AIM/phenotyping spectral flow cytometry panel is shown in Supplementary Table 2. To reduce bias associated with traditional gating techniques, we generated a Uniform Manifold Approximation and Projection (UMAP) of all collected CD4^+^ AIM^+^ T cells after dimensionality reduction and clustering. We identified 12 CD4^+^ AIM^+^ clusters across all study time points (Figure 2A). UMAPs split by timepoint are shown in Supplementary Figure 3A-B.

**Figure 2:**
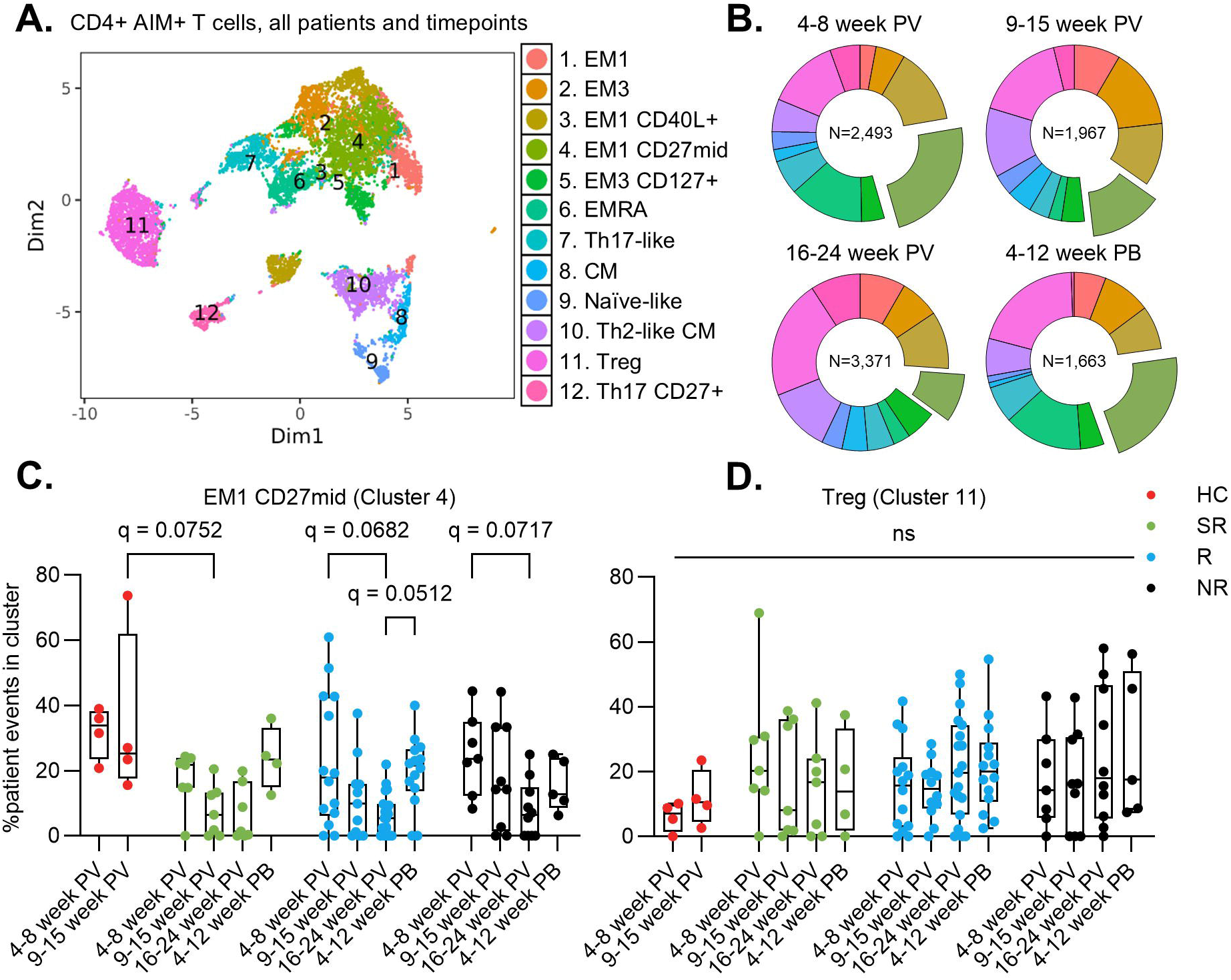
Long-term CD4^+^ AIM^+^ T cell memory responses to SARS-CoV-2 Spike preserved in OCR-treated pwMS. **(A)** UMAP depicting CD4^+^ AIM^+^ T cell (OX40^+^ CD137^+^) clusters from OCR-treated pwMS at all timepoints following spike peptide stimulation (left), with the corresponding cellular phenotype assigned to each cluster (right). Phenotypic clusters (right) include: T regulatory (Treg, CCR7-CD45RA-CD27^+^ FOXP3^+^ CD25^+^ CD127-), effector memory 1 (EM1, CCR7-CD45RA-CD27^+^), Th2 (CCR4^+^ CD45RA-CD27^+^), Th17 (CCR7-CD45RA-CD27^+^ CCR6^+^), EMRA (CCR7-CD27-CD45RA^+^ GZMB^+^), Naïve (CCR7^+^ CD45RA^+^ CD27^+^), central memory (CM, CCR7^+^ CD45RA-CD27^+^ CD25^+^), circulating T follicular helper (cTfh, CCR7^+^ CD45RA-CD27^+^ CXCR5^+^), Th1 (CCR7-CD45RA-CD27^+^ CXCR3^+^). **(B)** Proportional distribution of CD4^+^AIM^+^ clusters across post-vaccine time points, with total cell numbers indicated at the center of each circle graph. **(C-D)** Longitudinal analysis of (C) EM1 (cluster 4) and (D) Treg (cluster 11) in OCR-treated pwMS stratified by SR (green), R (blue), and NR (black) compared with healthy controls (red). Statistical testing was performed using Kruskal-Wallis test followed by the two-stage step-up method of Benjamini, Krieger, and Yekutieli. Q value is displayed. Q<0.05 (*). NS= not significant.

Effector memory (EM) T cells, particularly the EM1 subset, were the predominant population observed following SARS-CoV-2 vaccination (Figure 2B) (7). At 4 weeks post-vaccination, EM1 CD27^mid^ cells (cluster 4) expanded in pwMS and healthy controls, then contracted at 24 weeks (Figure 2C) (29). Expansion of EM1 CD27^mid^ cells after booster vaccination was reduced in the NR group compared with the R and SR groups. T regulatory (Treg) cells (cluster 11) increased over time from vaccination (Figure 2B), with no differences between response groups (Figure 2D). Taken together, these observations suggest that OCR treatment does not impair the magnitude or durability of antigen-specific CD4^+^ T cell responses following SARS-CoV-2 vaccination in pwMS.

### The SARS-CoV-2 Spike-specific CD8 T cell response is dominated by the effector memory (EMRA) phenotype

CD8^+^ AIM^+^ cells were defined as CD8^+^ T cells co-expressing CD69 and CD137, while the relative expression of additional surface and intracellular markers was used to assign phenotypes to identified clusters (Supplementary Figure 4). UMAPs split by post-vaccination time points are shown in Supplementary Figure 5A-B. Eight distinct CD8^+^ AIM^+^ clusters were identified, with five corresponding to EMRA phenotypes (Figure 3A). At 4 weeks post-vaccination, early memory CD25^+^ (cluster 2) were the dominant population, declining at later time points (Figure 3B). These cells were proportionally higher in the NR group at 4-8 weeks postvaccination (Supplementary Figure 5C). EM1 CD8^+^ T cells were observed across all groups, with no significant differences compared with healthy controls (Figure 3C). While not statistically significant, the SR group demonstrated the largest EM1 expansion following booster vaccination, suggesting that while robust T cell responses can occur in the absence of humoral immunity, coordinated humoral and cellular responses are more likely to confer optimal protection (Figure 3C) (30).

**Figure 3:**
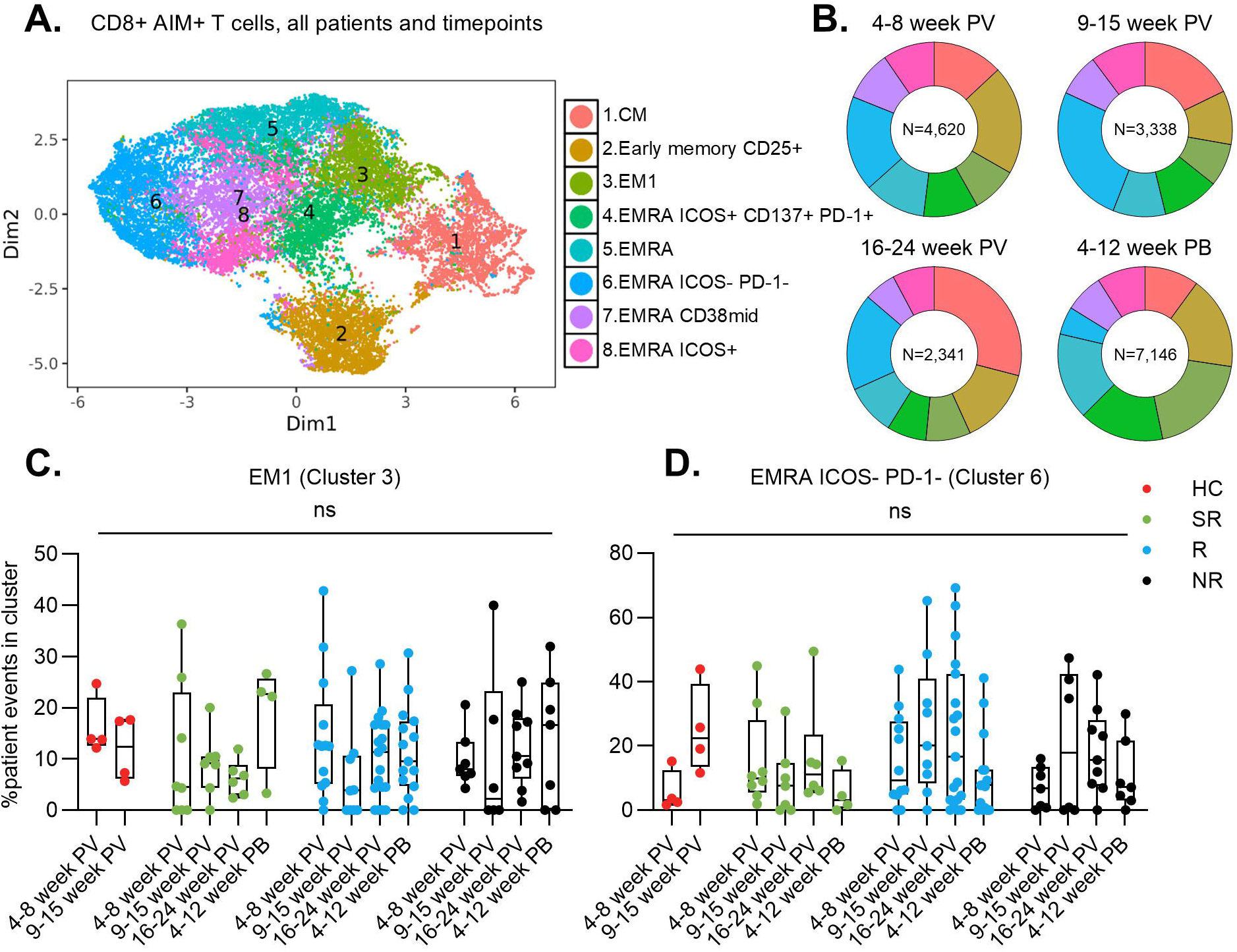
CD8^+^ AIM^+^ EMRA T cells dominate response to SARS-CoV-2 Spike protein in OCR-treated pwMS. **(A)** UMAP depicting CD8^+^ AIM^+^ T cell (CD69^+^ CD137^+^) clusters from OCR-treated pwMS at all timepoints following spike peptide stimulation (left), with the corresponding cellular phenotype assigned to each cluster (right). Phenotypic clusters (right) include: EMRA (CCR7-CD45RA^+^ CD27-GZMB^+^), EM1 (CCR7-CD27^+^ CD45RA-), EM3 (CCR7-CD27-CD45RA-), CM (CCR7^+^ CD45RA-CD27^+^ CD25^+^), Treg (CCR7-CD45RA-CD27^+^ FOXP3^+^ CD25^+^ CD127-), cTfh (CCR7^+^ CD45RA-CD27^+^ CXCR5^+^). **(B)** Proportional distribution of CD8^+^AIM^+^ clusters across post-vaccine time points, with total cell numbers indicated at the center of each circle graph. **(C-D**) Longitudinal analysis of (C) EM1 (cluster 3) and (D) EMRA (cluster 6) in OCR-treated pwMS stratified by SR (green), R (blue), and NR (black) compared with healthy controls (red). Statistical testing was performed using Kruskal-Wallis test followed by the two-stage step-up method of Benjamini, Krieger, and Yekutieli. Q value is displayed. Q<0.05 (*). NS= not significant.

The largest EMRA cluster (cluster 6), characterized by a lack of ICOS or PD-1 expression but expression of Granzyme B, peaked at 9-24 weeks post-vaccination, often more abundant in pwMS than in healthy controls (Figure 3D). Central memory (CM) CD8^+^ T cells were also detected, without group-specific differences (Supplementary Figure 5D). Collectively, these data support the notion that OCR therapy does not substantially alter the CD8^+^ T cell response to vaccination and highlight the prominent role of antigen-specific CD8^+^ T cells in the immune response to SARS-CoV-2 in OCR-treated pwMS.

### DN2 B cells represent a heterogeneous population more likely to persist following OCR-mediated B cell depletion

Next, we analyzed B cell repletion dynamics in peripheral blood following OCR infusion. To assess B cell reconstitution, dimensionality reduction and clustering were performed on CD19^+^ B cells, providing a comprehensive overview of the peripheral CD19^+^ B cell landscape in OCR-treated pwMS (Figure 4A). Given that OCR depletes most mature circulating CD20^+^ B cells, we hypothesized that residual B cells contribute to the observed heterogeneity in humoral immune responses (9). Following the observation that bulk T cell profiles remained stable over the post-infusion timeline, we then focused on B cell subpopulations capable of supporting humoral immune responses, including transitional, mature naïve, memory, and DN2-like memory B cells (Supplementary Figure 6A-C) (23, 26). From 18 identified CD19^+^ B cell clusters, feature plots were used to further visualize DN2 (IgD-CD27-CD11c^+^T-bet^+^) and memory B cell markers (Figure 4B). The heatmap used to assign cluster phenotypes is shown in Supplementary Figure 7A.

**Figure 4:**
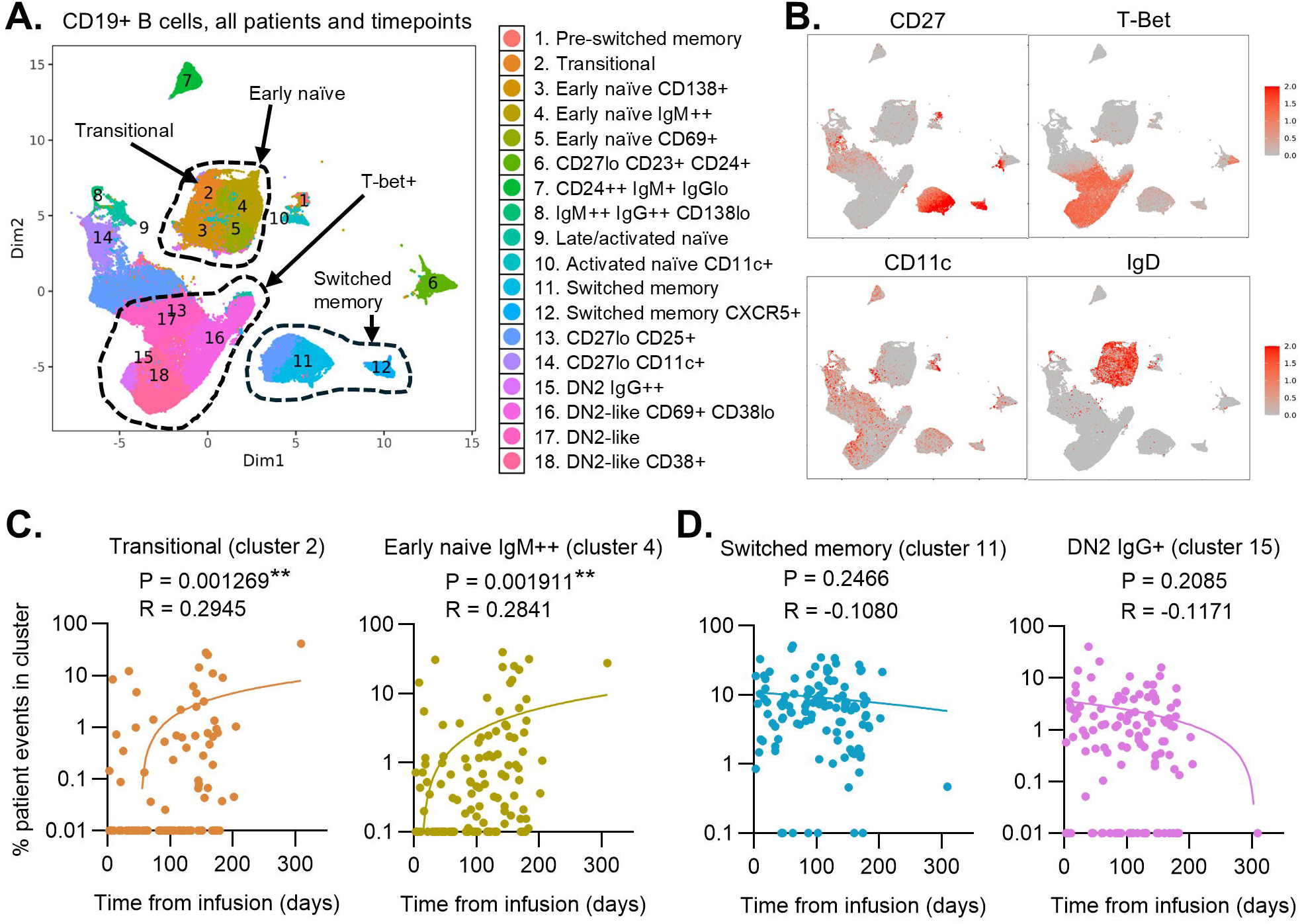
Switched memory and DN2 B cells are resistant to OCR-mediated B cell depletion. **(A)** UMAP projection of all CD19^+^ B cells from OCR-treated pwMS (left) with cluster identities indicated (right). **(B)** Feature plots of lineage-defining markers (CD27, T-bet, CD11c, and IgD). **(C-D)** Longitudinal kinetics of B cell subsets, plotted as correlations between the relative proportions of transitional (C, left), naïve (C, right), switched memory (D, left), and DN2 (D, right) B cell clusters and days from OCR infusion. Each dot represents the percentage of CD19^+^ cells from one patient within that specific cluster, out of the total number of CD19^+^ cells from that patient. Spearman’s rank correlation coefficient was used to assign statistical significance. p<0.05 (*), p<0.01 (**).

Transitional (cluster 2) and early naïve (cluster 4) B cells showed positive correlations with time from infusion, indicating both susceptibility to depletion and effective peripheral repletion in peripheral blood (Figure 4C). In contrast, switched memory (cluster 11) and DN2 IgG^+^ (cluster 15) B cells did not exhibit this repletion pattern, instead showing non-significant negative correlations with days from infusion (Figure 4D). All four identified DN2-like B cell clusters (clusters 15-18) remained stable over time following infusion, consistent with relative resistance to depletion (Figure 4D, Supplementary Figure 7B).

### Spectral flow cytometry reveals persistence of DN2 and classical memory B cells despite OCR infusion

A manual gating strategy was employed to confirm the clustering results (Figure 5A) (26). Total CD19^+^ B cells showed a positive but non-significant correlation with time from infusion (Figure 5B). Transitional and mature naïve B cells demonstrated significant repletion, whereas DN2 B cells did not exhibit such repletion kinetics, indicating relative resistance to OCR-mediated depletion (Figure 5C, D). Thus, both unsupervised clustering and manual gating approaches converge on the conclusion that certain ASC-precursor populations, including DN2 B cells, may resist OCR-mediated B cell depletion and serve as the cellular substrate for humoral responses in a subset of pwMS treated with OCR.

**Figure 5:**
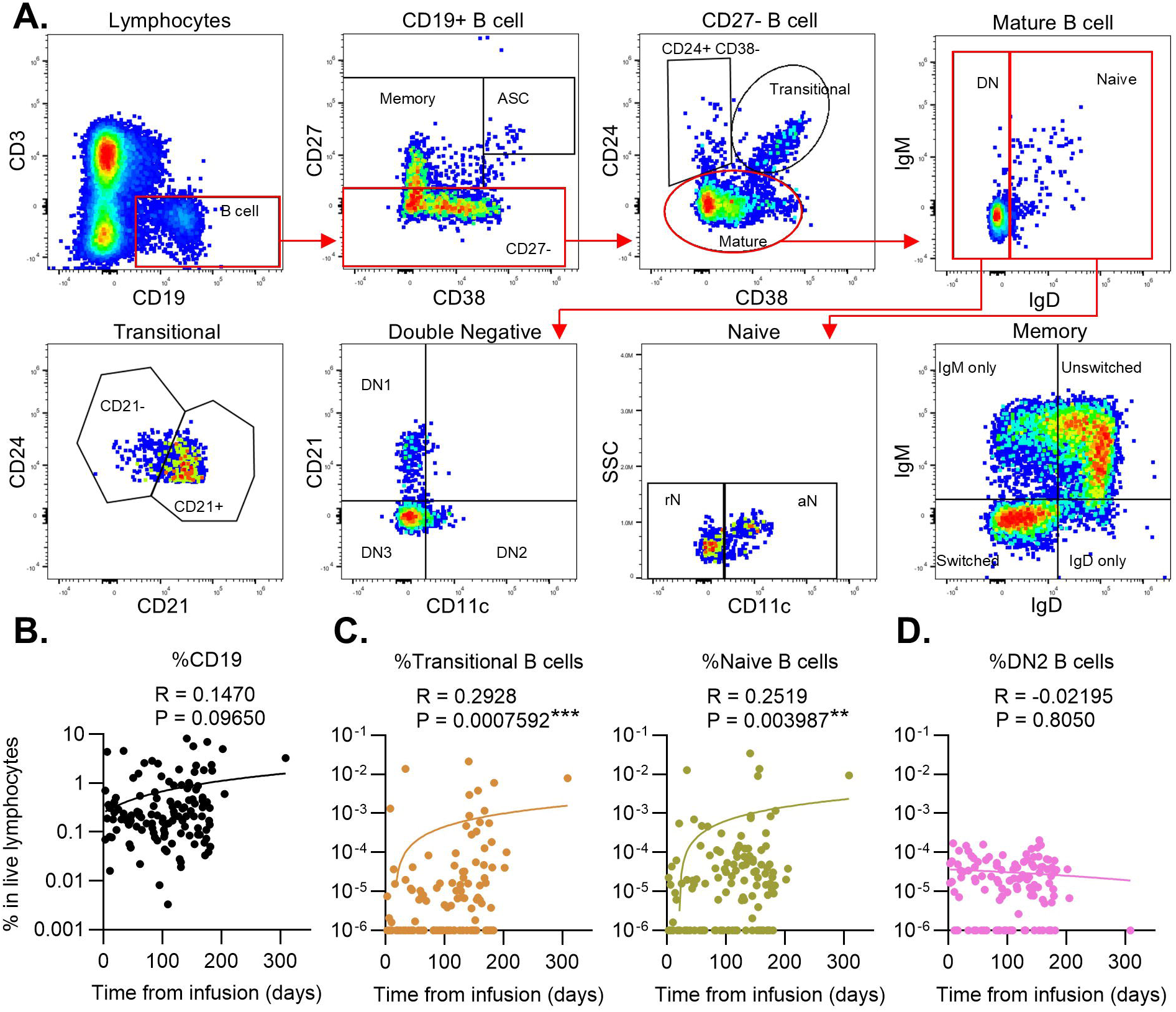
Peripheral repletion of CD19^+^ early B cell subsets differs from that of total CD19^+^ B cells. **(A)** Gating strategy for identifying B cell subsets from the total live lymphocyte population. **(B)** Correlation of total CD19^+^ B cells (% of live lymphocytes) and days from OCR infusion **(C)** Correlations demonstrating subset-specific repletion kinetics following OCR infusion for transitional (left) and naïve (right) B cells, expressed as % of live lymphocytes. **(D)** Correlation of DN2 B cells (% of live lymphocytes) and days from OCR infusion. Spearman’s rank correlation coefficient was used to assign statistical significance. p<0.05 (*), p<0.01 (**), p<0.001 (***).

### DN2 and mature naive B cells are enriched in SR compared to NR in OCR-treated pwMS

The accurate identification of DN2 B cells among other DN B cell subtypes was validated using histograms showing DN2-specific marker expression (Figure 6A) (23, 25). In the SR group, the total CD19^+^ B cell population increased with time from infusion, whereas the opposite trend was observed in the NR group (Figure 6B). Transitional and naïve B cells demonstrated more robust repletion in the SR group compared with NR, with significant differences observed for naïve B cells (Figure 6B). DN2 B cells appeared relatively resistant to OCR-mediated B cell depletion in both groups (Figure 6B). The observed consistent repletion of peripheral B cell subsets, as well as total CD19^+^ B cells, in SR patients compared to the NR patients suggests the regulation of B cell reconstitution, and subsequent differentiation of repopulating subsets, may determine the capacity to mount a vaccine-induced humoral response.

**Figure 6:**
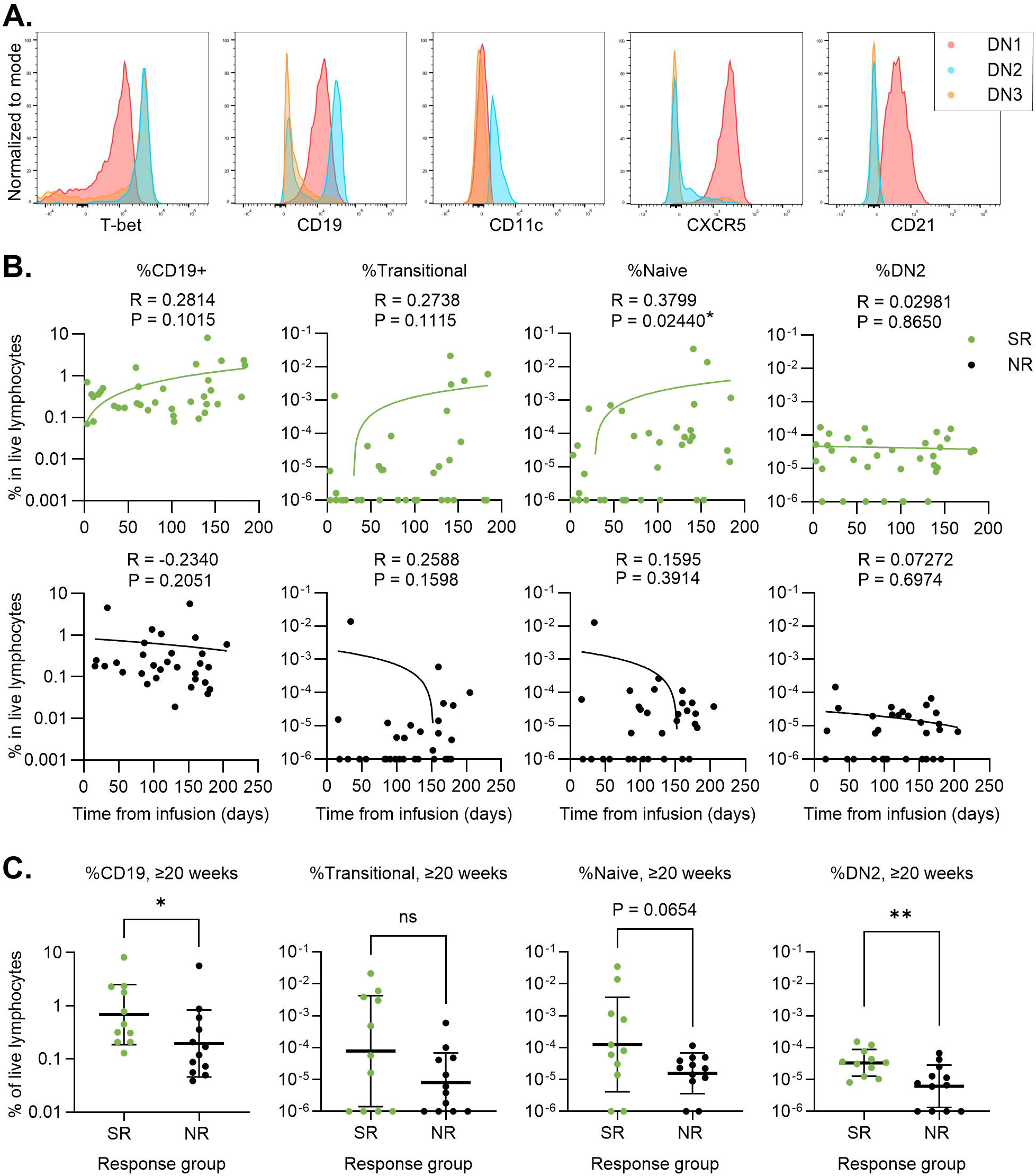
Naïve and DN2 B cell subsets are associated with robust humoral immune responses to SARS-CoV-2 vaccination. **(A)** Representative histograms depicting flow cytometry DN subsets, defined as DN1 (red), DN2 (blue), and DN3 (yellow), based on the normalized MFI of 5 established lineage markers: T-bet, CD19, CD11c, CXCR5, and CD21. **(B)** Peripheral reconstitution of total B cell and B cell subsets compared between SR (top row, green) and NR (bottom row, black) OCR-treated pwMS. Correlations depict repletion kinetics following OCR infusion for (from left to right) total CD19^+^, transitional, naïve, and DN2 B cells, expressed as % of live lymphocytes. **(C)** Comparison of (from left to right) total CD19^+^, transitional, naïve, and DN2 B cells (expressed as % of live lymphocytes) between OCR-treated pwMS from the SR and NR groups, including only samples collected at ≥20 weeks from most recent OCR infusion. Spearman’s rank correlation coefficient or Mann-Whitney U test was used to assign statistical significance. P<0.05 (*), p<0.01 (**). NS= not significant.

At ≥20 weeks post-infusion, the SR group exhibited a higher proportion of CD19^+^ B cells than the NR group, supporting the concept that impaired or dysregulated B cell repletion in NR patients results in reduced overall B cell recovery, particularly at later time points (Figure 6C). Transitional, mature naïve, and DN2 B cells were also more abundant in the SR group, with the most significant difference observed in the DN2 subset (Figure 6C). Collectively, these findings support a model in which transitional, naïve, and OCR-resistant DN2-like B cells serve as key precursors of vaccine-induced anti-Spike IgG in pwMS treated with OCR, potentially underlying the heterogeneity in humoral vaccine responses observed in these patients.

## Discussion

OCR, an anti-CD20 antibody-based B-cell-depleting therapy, benefits pwMS by effectively reducing disease burden and slowing disability progression. However, this therapeutic effect may impair *de novo* humoral immune responses, increasing susceptibility to COVID-19 infection and diminishing the protective efficacy of vaccinations (2, 3). Prior studies investigating immune responses in OCR-treated pwMS have varied in emphasis, including B-cell repopulation kinetics during different OCR infusion regimens and antigen-specific cellular responses to SARS-CoV-2 vaccination (4, 7, 18, 31, 32). Our group, as well as others, has demonstrated that while T-cell responses to SARS-CoV-2 vaccination and infection are largely preserved under OCR, only a minority of pwMS retain the capacity to mount robust humoral responses (8, 12). Among the main objectives of this study was to investigate the cellular correlates of the preserved humoral immune response observed in a subset of pwMS treated with OCR. Specifically, we sought to determine whether qualitative features of immune reconstitution, rather than total circulating B-cell counts alone, distinguish patients who can mount robust vaccine-induced antibody responses.

Based on post-vaccination Spike IgG levels, we defined three response groups: SR, R, and NR, which were compared throughout the study. Consistent with previous reports, antigen-specific CD4^+^ AIM^+^ EM1 T cells were predominantly preserved in OCR-treated pwMS and contracted between 4 and 24 weeks after vaccination, providing evidence of long-term CD4^+^ memory responses comparable to healthy controls (7, 29). The more robust CD4^+^ AIM^+^ EM1 response in the SR and R groups compared to the NR group underscores that while humoral and cellular responses to SARS-CoV-2 vaccination are partially decoupled in OCR-treated pwMS, OCR therapy does not preclude coordination between these arms of the immune response (8, 12, 30). In our cohort, longitudinal changes in antigen-specific CD4^+^ cTfh cells were not observed, although this may be explained by previously described vaccine-induced shifts in this cell population occurring prior to patient sample collection at 4 weeks post-vaccination (7). Among the CD8^+^ AIM^+^ T cells, the NR group showed increased antigen-specific EMRA responses, potentially representing a compensatory expansion of cytotoxic CD8^+^ T cells in the absence of humoral immune responses as reported previously (7). In line with this, antigen-specific CD8^+^ early memory cells were proportionally greater in NR patients. The increased AIM^+^ CD8^+^ T cell subtypes in the NR group suggest that the compensatory CD8^+^ EM1 T cell activation previously reported in OCR-treated pwMS lacking an effective humoral immune response to SARS-CoV-2 vaccination may extend to additional CD8^+^ memory T cell subtypes. The observed heterogeneity in T cell responses among OCR-treated pwMS, regardless of humoral response group, may also relate to the varied haplotypes present in the patient population (33). While it has been shown that a subset of inflammatory CD8^+^ CD20^+^ memory T cells is selectively depleted by OCR treatment, our patients had already been receiving OCR infusions for several years, limiting our ability to evaluate early treatment-related immune shifts that may partially explain reduced virus-specific T cell responses (34–36). These observations extend prior findings regarding the SARS-CoV-2 antigen-specific T cell response by incorporating longitudinal data and additional humoral responder-group comparisons, highlighting the interplay between cellular and humoral immunity in OCR-treated pwMS. Together, these findings reinforce the concept that anti-CD20 therapy attenuates antibody responses more than T-cell immunity, although coordinated responses remain detectable in a subset of patients.

Studies investigating extended-interval dosing (EID) of OCR have not observed an association with increased risk of disease relapses or disease progression in pwMS (32, 37, 38). EID enables more robust B-cell reconstitution before subsequent infusions, potentially enhancing the capacity for humoral immune responses to vaccination. It is therefore critical to identify cellular subsets that may underlie humoral immune responses to vaccination in OCR-treated pwMS. Our study provides mechanistic insights into the heterogeneity of SARS-CoV-2 spike-specific antibody responses. As expected, transitional and mature naïve B cells replenished in a time-dependent manner following infusion, consistent with prior studies (32, 39). Faster repletion of naïve B cells in SR patients suggests a replenished precursor pool available for antigen-specific activation, potentially enhancing memory formation and humoral immunity (14). However, this contrasted with the observation that DN2 and a subset of switched memory B cells were relatively resistant to OCR-mediated depletion, remaining detectable even at early post-infusion time points, as observed using two independent methods of analysis. This is in line with the previously observed persistence of activated memory B cells following OCR therapy (40). DN2-like B cells, identified by T-bet and CD11c expression in the absence of CXCR5 and CD21, represent a phenotypically heterogeneous subset that has been previously implicated in extrafollicular antibody responses, including in the context of SARS-CoV-2 vaccine responses in healthy subjects (23, 25, 26, 41). These cells may additionally serve as precursors to ASCs (42, 43). In line with this, we observed that OCR-treated pwMS with greater post-vaccination Spike-IgG production exhibited not only a significant enrichment of the DN2 B cell population but also a more robust and effective repletion of transitional and mature naïve B cell subsets in the periphery. These results suggest that some infusion-resistant DN2 and memory subsets, combined with the accelerated recovery of early B cell populations, may underlie the observed greater humoral response capacity of some OCR-treated pwMS. Importantly, these findings suggest that total peripheral CD19^+^ B-cell counts may incompletely capture immune competence after OCR, as the composition of the recovering B-cell compartment appears more informative than absolute B-cell numbers alone. Clinically, the abundance of DN2-like B cells and the overall dynamics of B cell reconstitution (particularly of mature naïve B cells) may serve as biomarkers of vaccine responsiveness, although further studies are required to validate these observations. If confirmed prospectively, such biomarkers could help individualize vaccine timing, prioritize booster doses, or identify patients who may benefit from alternative aCD20 dosing strategies.

Our study has limitations. The delineation of pwMS into SR, R, and NR groups further limited the sample size and complicated statistical comparisons. To partially address this, we incorporated data from a parallel cross-sectional study that used harmonized protocols, thereby increasing statistical power and generalizability. Variability in prior OCR therapy duration across patients is another potential confounder. However, we found that the recurring OCR-induced depletion and repletion dynamics were not significantly affected by the total number of previous infusions. By stratifying patients by the latest infusion time, we controlled for time since infusion. Finally, while our phenotypic data suggest that DN2 and switched memory subsets correlate with humoral responsiveness following SARS-CoV-2 vaccination, direct functional validation linking these populations to antibody-secreting activity remains ongoing. In addition, because this was a single-center cohort studied during a specific vaccine era, broader validation across populations, vaccine platforms, and infectious exposures will be important.

In summary, our study provides mechanistic insight into the immunologic heterogeneity of vaccine responses in OCR-treated pwMS. We demonstrate that SR patients are distinguished not only by accelerated naïve B cell repletion but also by the persistence and enrichment of DN2 and some memory B cell subsets following OCR-mediated B cell depletion, which may fuel antibody responses via extrafollicular and classical memory pathways. These findings identify potential biomarkers that may predict humoral responsiveness to vaccination and thereby contribute to higher anti-Spike IgG titers in this patient population. Clinically, quantifying these subsets may offer greater predictive utility than total CD19^+^ counts and could guide personalized immunization strategies in pwMS receiving OCR. More broadly, our results highlight the need to consider infusion-resistant memory compartments when evaluating immune competence under B-cell-depleting therapies. These observations may also be relevant beyond SARS-CoV-2 vaccination, including for other vaccines and for future emerging pathogens that require effective *de novo* humoral immunity.

## Methods

### Patients and Samples

Patient demographics are detailed in Supplementary Table 1. This prospective, observational study included 30 patients enrolled in the Vaccine generated Immunity in Ocrelizumab-treated patients: Longitudinal Assessments (VIOLA) study (NCT04843774), which incorporated predefined time points: baseline (pre-vaccination), 4, 12, and 24 weeks post-vaccine, and 4 and 12 weeks post-booster vaccination, as well as 36 pwMS from a predominantly cross-sectional study (NCT04682548), for a total of 66 pwMS treated with ocrelizumab (8, 12). Serum and peripheral blood mononuclear cells (PBMCs) were isolated from each patient sample and cryopreserved in liquid nitrogen. Longitudinal healthy controls were obtained from the NYU Vaccine Center and included PMBCs collected from baseline through 12 weeks post-vaccination and serum samples through 24 weeks post-vaccination.

### Analysis of SARS-CoV-2 Spike-specific IgG in patient sera

A proprietary multiplex bead-based immunoassay was used to quantify anti-SARS-CoV-2 IgG, as previously described (8, 12, 44). Briefly, fluorescent barcoded magnetic beads (MagPlex microspheres; Luminex) were coupled to the Wuhan variant full-length spike protein (Sino Biological cat no. 40590-V08B). Beads conjugated to tetanus toxoid, human serum albumin, and anti-human IgG were also included as in-assay controls (Jackson Immunoresearch). Binding was detected using the MagPix platform (Luminex).

### Analysis of ocrelizumab concentration in MS patient sera

Serum OCR concentrations were measured using a proprietary anti-ocrelizumab enzyme-linked immunosorbent assay (ELISA) (Sanquin Diagnostic Services, Amsterdam, The Netherlands).

### SARS-CoV-2 Spike Peptide Library

PBMCs were stimulated using a 15-mer overlapping peptide pool spanning the SARS-CoV-2 spike glycoprotein (11-amino acid overlap; Miltenyi Biotec PepTivator ® SARS-CoV-2 Prot_S Complete, cat. 130-129-712).

### SPECTRAL FLOW CYTOMETRY

#### Antibody panel

A 35-parameter spectral flow cytometry panel was designed for the Cytek Aurora Spectral Analyzer (Cytek Biosciences). Panel details are provided in Supplementary Table 3.

#### Sample staining and acquisition

PBMC (∼1×10^6^ cells per sample) were thawed, washed, and resuspended in FACS buffer (Hanks balanced salt solution supplemented with 2% fetal bovine serum). Fc receptor blocking was performed using Human TruStain FcX and True-Stain Monocyte Blocker (BioLegend cat. Nos. 422302 and 426102). Surface staining was performed for 30 min at room temperature (22 °C) in a final volume of 50 μL, followed by washing and viability staining with the Zombie NIR™ Fixable Viability Kit (BioLegend cat. 423106). Cells were then fixed and permeabilized using the eBioscience™ Foxp3 / Transcription Factor Staining Buffer Set (ThermoFisher Scientific, cat. 00-5523-00), followed by intracellular staining for 1 hour at 4 °C. Data were acquired on the Cytek Aurora Spectral Analyzer using SpectroFlo software. An unstimulated condition was included for each sample to control for background activation.

#### Initial analysis and quality control

FlowJo v10 (BD Biosciences) was used to gate CD19^+^ B cells and both CD4^+^ and CD8^+^ activation induced marker-positive (AIM^+^) T cells. Exported CSV files were further analyzed in R, and samples with cell viability <50% were excluded. Additionally, patient samples with < 5 AIM^+^ T cells were excluded from T cell analysis but retained for B cell analysis due to the challenges of accurately gating cell populations of this size. Graphs were generated using GraphPad Prism (v10.3.0).

#### Bioinformatics processing of spectral flow cytometry data

Marker intensity data were exported from FlowJo and normalized within each batch using Mutual Nearest Neighbors (MNN) correction (45). Data were randomly down-sampled to one-third, aligned, and clustered using the Louvain algorithm. Dimensionality reduction was performed using Uniform Manifold Approximation and Projection (UMAP) and t-distributed Stochastic Neighbor Embedding (t-SNE). Cluster identities were manually annotated based on marker expression patterns and heatmap visualization. Analyses were conducted in R (v4.5.0) with ggplot2 (v3.4.1).

## Statistical Analysis

All statistical analyses were conducted using GraphPad Prism (v9.0.0/v10.3.0). For intra-group and inter-group comparisons (Figure 1B), a mixed-effects model with the Geisser-Greenhouse correction followed by a post-hoc Tukey’s Honest Significant Difference (HSD) test was used to correct for false discovery rate (FDR). For comparisons among three groups, the Kruskal-Wallis test followed by Dunn’s post-hoc test was applied. For comparisons between two groups, the Mann-Whitney U test was used.

Correlations were assessed using Spearman’s rank correlation coefficient. For CD4^+^ AIM^+^ and CD8^+^ AIM^+^ datasets, FDR correction was performed using the two-stage step-up method of Benjamini, Krieger, and Yekutieli, given the large number of comparisons and small sample sizes. Adjusted q values are reported for these analyses, with q <.05 considered statistically significant. For all other analyses, p <.05 was considered significant (*), with additional thresholds: p<.01 (**), p<.001 (***), and p<.0001 (****). Non-significant results are denoted “NS”.

## Standard protocol approvals, registrations, and patient consents

Written informed consent was obtained from all participants prior to enrollment and sample collection. All procedures were conducted under the approved IRB protocols s21-00206 for the VIOLA study [NCT04843774], and s20-01799 for the cross-sectional study [NCT04682548].

## Data availability

All anonymized data not published within this article will be made available upon request from qualified investigators. The principal author takes full responsibility for the data, the analysis and interpretation, and the conduct of the research. The principal author has full access to all data, and the right to publish all data separately from the guidance of the sponsor.

## Acknowledgements

The authors thank Ramin Herati (NYU Grossman School of Medicine), for his guidance and assistance in the design of the 35-color spectral flow cytometry panel and statistical analysis of data. We also thank Matthew Lustberg, Laura Alice, and Tom Walsh for their assistance with the financial and administrative management of this project, as well as the Krogsgaard lab members for processing and storing patient blood samples and providing technical support.

## Supplementary Figure Legends

**Supplementary Table 1: MS patient demographics.** PwMS are grouped by response category, from left to right: super-responders (green), responders (blue), and non-responders (gray). All values were obtained from each patient’s baseline study visit. Variables include sex, race, ethnicity, MS subtype, vaccine type, and prior SARS-CoV-2 infection before baseline sample collection. Variables compared across the three patient response groups included age, body mass index (BMI), MS disease duration, time from ocrelizumab (OCR) infusion to vaccination, time from OCR infusion to baseline collection, total time on OCR at baseline collection, and average interval between infusions. The Kruskal-Wallis test was used to determine statistical significance. p<0.05 (*) was considered significant.

**Supplementary Figure 1: OCR infusion history and demographics of pwMS response groups. (A)** Comparison of %CD19^+^ B cells in OCR-treated pwMS at baseline between the SR (green), R (blue), and NR (black) response groups. **(B)** Comparison of OCR treatment history and patient demographics in OCR-treated pwMS between SR (green), R (blue), and NR (black). Variables compared include (from left to right): MS disease duration, days from OCR infusion to baseline sample collection, patient age, and the average number of days between OCR infusions. The Kruskal-Wallis test followed by Dunn’s post-hoc test was used to determine statistical significance. NS= not significant.

**Supplementary Table 2: Antibody panel used for spectral flow cytometry analysis.** Column headers (left to right): marker (cellular marker targeted by antibody), fluorophore (conjugated fluorophore used for detection), and manufacturer.

**Supplementary Figure 2: Heatmap used for phenotyping CD4^+^ AIM^+^ cells.** Heatmap displaying T cell markers used to phenotype CD4^+^ AIM^+^ clusters. Each cellular marker is denoted on the right side, with the top numbers indicating cluster ID. The median expression of each cellular marker is measured as median fluorescence intensity (MFI).

**Supplementary Figure 3: Distribution of CD4^+^ AIM^+^ clusters in healthy controls. (A)** Representative UMAPs of CD4^+^ AIM^+^ T cells (OX40^+^CD137^+^) from OCR-treated pwMS following spike peptide stimulation, separated by post-vaccination timepoints. The phenotype assigned to each cluster is shown on the right. **(B)** Representative UMAPs of CD4^+^ AIM^+^ T cells from healthy controls (HC), divided by post-vaccination timepoints.

**Supplementary Figure 4: Heatmap used for phenotyping of CD8^+^ AIM^+^ cells.** Heatmap displaying T cell markers for the phenotyping of CD8^+^ AIM^+^ clusters. Each cellular marker is denoted on the right side, with the top numbers indicating cluster ID. The median expression level of each cellular marker is measured and reported as MFI.

**Supplementary Figure 5: Longitudinal analysis of CD8^+^AIM^+^ T cells. (A)** Representative UMAPs of CD8^+^ AIM^+^ T cells (OX40^+^CD69^+^) from OCR-treated pwMS following spike peptide stimulation, split by post-vaccination timepoints. The phenotype assigned to each cluster is shown on the right. **(B)** Representative UMAPs of CD8^+^ AIM^+^ T cells from HC, split by post-vaccination timepoints. **(C-D)** Longitudinal trajectories of CD8^+^ AIM^+^ T cell phenotypes compared between OCR-treated pwMS stratified by SR (green), R (blue), and NR (black), as well as in HC (red). Clusters analyzed included (C) early memory (cluster 2) and (D) activated central memory (CM, cluster 1). Statistical significance was determined using the Kruskal–Wallis test, followed by the two-stage step-up method of Benjamini, Krieger, and Yekutieli. Q value is displayed. Q<0.05 (*). NS= not significant.

**Supplementary Figure 6: Bulk T cell phenotypes are unaffected by time since OCR infusion. (A)** Correlations demonstrating repletion kinetics following OCR infusion of bulk CD4^+^ T helper subsets, expressed as a percentage of all non-naïve (nn) CD4^+^ T cells. Subsets analyzed include (from left to right): cTfh, Th1/Th17, Th1 CCR4-, Th17, and Th2 CCR4^+^. (B) Correlations demonstrating repletion kinetics following OCR infusion of bulk CD4^+^ memory subsets, expressed as a percentage of all CD4^+^ T cells. Subsets analyzed include (from left to right): EMRA, CM, EM1, EM2, and EM3. (C) Correlations demonstrating repletion kinetics following OCR infusion of bulk CD8^+^ memory subsets, expressed as a percentage of all CD8^+^ T cells. Subsets analyzed include (from left to right): EMRA, CM, EM1, EM2, and EM3. Spearman’s rank correlation coefficient was used to assign statistical significance. p<0.05 (*).

**Supplementary Figure 7: Heatmap of CD19^+^ B cells used for phenotyping B cell clusters. (A)** The heatmap displays CD19^+^ B cell markers used to define CD19^+^ B cell clusters from OCR-treated pwMS. Each cellular marker is denoted on the right side, with the top numbers indicating cluster ID. The median expression level of each cellular marker is measured and reported as MFI. **(B**) Correlation of (from left to right): DN2-like CD69^+^ CD38lo (cluster 16), DN2-like (cluster 17), and DN2-like CD38^+^ (cluster 18), and days from OCR infusion. Each dot represents the percentage of CD19^+^ cells from one patient within that specific cluster, out of the total number of CD19^+^ cells from that patient. Spearman’s rank correlation coefficient was used to assign statistical significance. p<0.05 (*).

