## Supplementary Figures for "Atypical memory B cells contribute to heterogeneous vaccine responses in ocrelizumab-treated patients with multiple sclerosis"

| | Super Responders (n=16) | | | | | Responders (n=34) | | | | | Non-Responders (n=16) | | | | | |
|---|---|---|---|---|---|---|---|---|---|---|---|---|---|---|---|---|
| **Variable** | ***N*** | **Percent** | **Mean** | **SD** | **Range** | ***N*** | **Percent** | **Mean** | **SD** | **Range** | ***N*** | **Percent** | **Mean** | **SD** | **Range** | **p-value** |
| Sex | | | | | | | | | | | | | | | | |
| Female | 12 | 75.0 | | | | 23 | 67.6 | | | | 11 | 68.8 | | | | |
| Male | 4 | 25.0 | | | | 11 | 32.4 | | | | 5 | 31.3 | | | | |
| Race | | | | | | | | | | | | | | | | |
| White | 2 | 12.5 | | | | 14 | 41.2 | | | | 8 | 50.0 | | | | |
| Black/African American | 7 | 43.8 | | | | 6 | 17.6 | | | | 4 | 25.0 | | | | |
| Other | 7 | 43.8 | | | | 14 | 41.2 | | | | 4 | 25.0 | | | | |
| Ethnicity | | | | | | | | | | | | | | | | |
| Hispanic/Latino | 6 | 37.5 | | | | 10 | 29.4 | | | | 5 | 31.3 | | | | |
| Not Hispanic/Latino | 10 | 62.5 | | | | 24 | 70.6 | | | | 11 | 68.8 | | | | |
| MS Subtype | | | | | | | | | | | | | | | | |
| RRMS | 16 | 100.0 | | | | 34 | 100.0 | | | | 16 | 100.0 | | | | |
| Vaccine Type | | | | | | | | | | | | | | | | |
| Pfizer | 9 | 56.3 | | | | 27 | 79.4 | | | | 10 | 62.5 | | | | |
| Moderna | 6 | 37.5 | | | | 7 | 20.6 | | | | 4 | 25.0 | | | | |
| Johnson and Johnson | 1 | 6.3 | | | | 0 | 0.0 | | | | 2 | 12.5 | | | | |
| Infection before collection | | | | | | | | | | | | | | | | |
| Yes | 1 | 6.3 | | | | 2 | 5.9 | | | | 0 | 0.0 | | | | |
| No | 15 | 93.8 | | | | 32 | 94.1 | | | | 16 | 100.0 | | | | |
| Age, years | | | 35.6 | 7.4 | 26.0 - 50.0 | | | 39.9 | 10.1 | 23.0 - 58.0 | | | 38.1 | 10.2 | 20.0 - 56.0 | 0.3658 |
| Body Mass Index | | | 34.0 | 9.6 | 19.3 - 54.3 | | | 28.4 | 8.6 | 19.2 - 57.6 | | | 27.5 | 7.1 | 17.2 - 46.1 | 0.0507 |
| Disease Duration, years | | | 8.1 | 5.5 | 1.6 - 20.1 | | | 9.1 | 8.8 | 1.6 - 34.7 | | | 12.6 | 8.9 | 2.6 - 35.0 | 0.3678 |
| Infusion to vaccination time, days | | | 129.3 | 51.6 | 48.0 - 197.0 | | | 117.2 | 55.0 | 33.0 - 233.0 | | | 77.1 | 51.0 | 21.0 - 190.0 | 0.0164* |
| Infusion to baseline collection time, days | | | 91.4 | 46.9 | 18.0 - 180.0 | | | 119.0 | 67.0 | 14.0 - 309.0 | | | 132.8 | 45.5 | 34.0 - 205.0 | 0.136 |
| Time on ocrelizumab at baseline, years | | | 1.9 | 1.1 | 0.4 - 4.0 | | | 2.3 | 1.2 | 0.2 - 4.2 | | | 2.7 | 1.1 | 0.5 - 4.2 | 0.0973 |
| Average time between infusions, days | | | 197.3 | 33.1 | 148.5 - 268.0 | | | 201.5 | 30.3 | 182.0 - 327.0 | | | 220.3 | 46.4 | 176.7 - 342.5 | 0.1343 |

### Supplementary Table 1

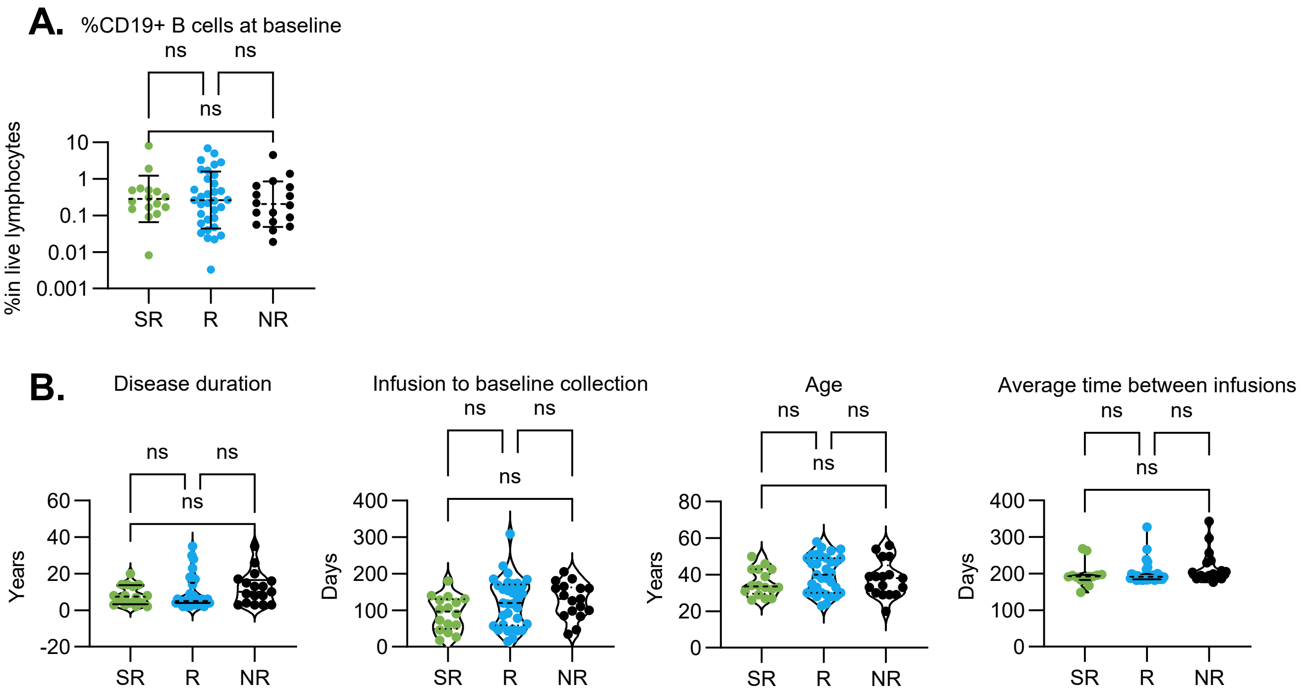


**Supplementary Figure 1**

| Marker | Fluorophore | Manufacturer |
|---|---|---|
| Live/Dead | LiveDead Blue | Fisher |
| CD3 | SparkBlue 550 | BioLegend |
| CD4 | SparkViolet 538 | BioLegend |
| CD8 | PE-Fire 640 | BioLegend |
| CD11c | PerCP | BioLegend |
| CD134 (OX40) | BUV805 | BD Biosciences |
| CD24 | BV480 | BD Biosciences |
| CD19 | BUV496 | BD Biosciences |
| CD137 (4-1BB) | A647 | BioLegend |
| CD21 | PE-Cy5 | BD Biosciences |
| CD23 | BUV615 | BD Biosciences |
| CD25 | BUV563 | BD Biosciences |
| CD27 | SB702 | Invitrogen |
| CD38 | APC/Fire810 | BioLegend |
| CD40 | PacBlue | BioLegend |
| CD45RA | Spark NIR 685 | BioLegend |
| IgM | BV570 | BioLegend |
| CCR6 | SB780 | Invitrogen |
| CD69 | BV650 | BioLegend |
| CD127 | SparkYG 581 | BioLegend |
| CD138 | BV510 | BioLegend |
| CD95 | BV421 | BioLegend |
| CD183 (CXCR3) | BV750 | BD Biosciences |
| CD185 (CXCR5) | BB515 | BD Biosciences |
| CD197 (CCR7) | BV605 | BioLegend |
| CD278 (ICOS) | APC-Fire750 | BioLegend |
| CD279 (PD-1) | BB700 | BD Biosciences |
| HLA-DR | BUV661 | BD Biosciences |
| IgD | BUV737 | BD Biosciences |
| Foxp3 | PE-Cy5.5 | Invitrogen |
| Tbet | PE-Cy7 | BioLegend |
| CD40L | PE-eF610 | Invitrogen |
| GzmB | A700 | BD Biosciences |
| CD194 (CCR4) | BUV395 | BD Biosciences |
| IgG (extracellular) | PerCP-Vio700 | Miltenyi |

**Supplementary Table 2**

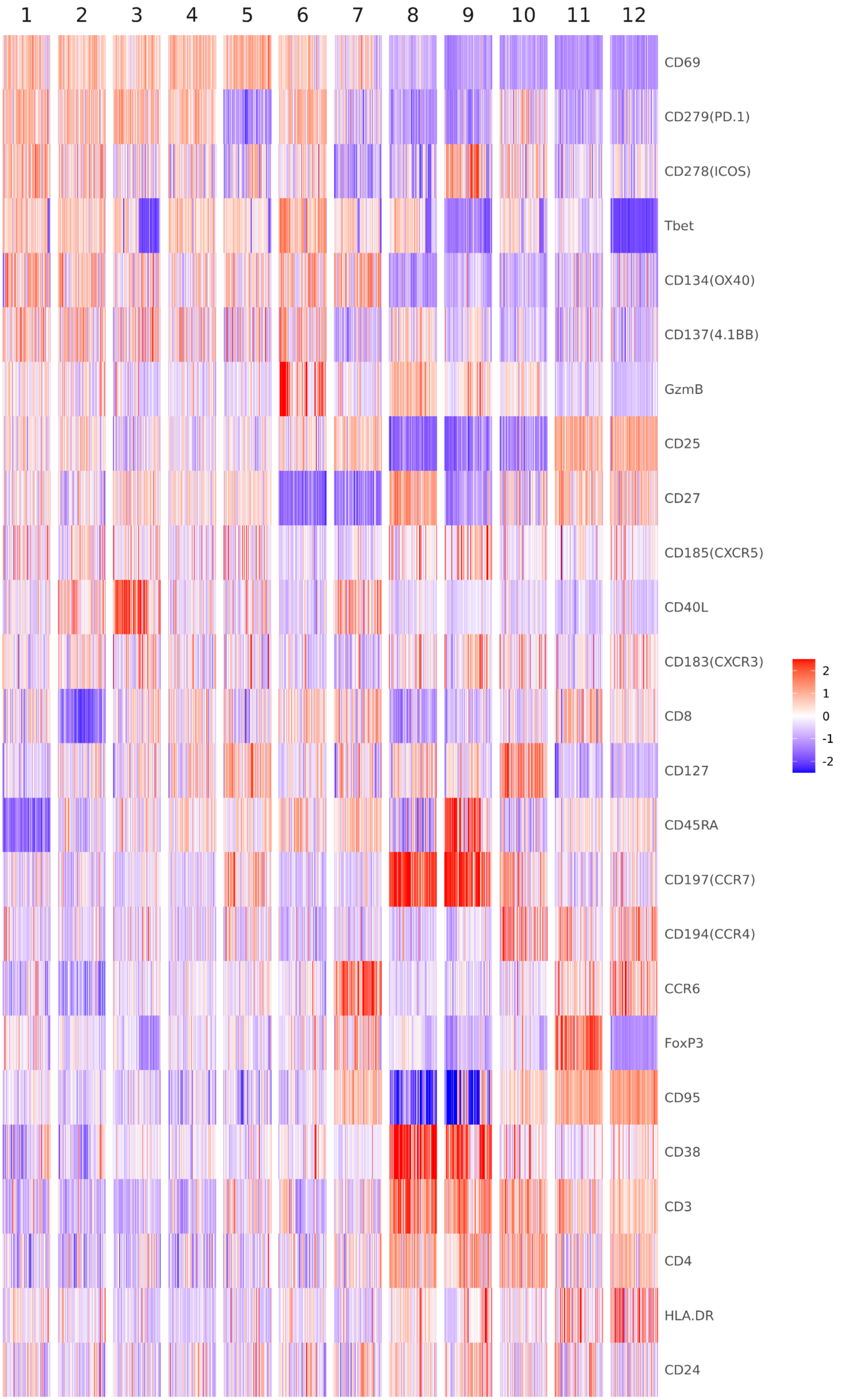


**Supplementary Figure 2**

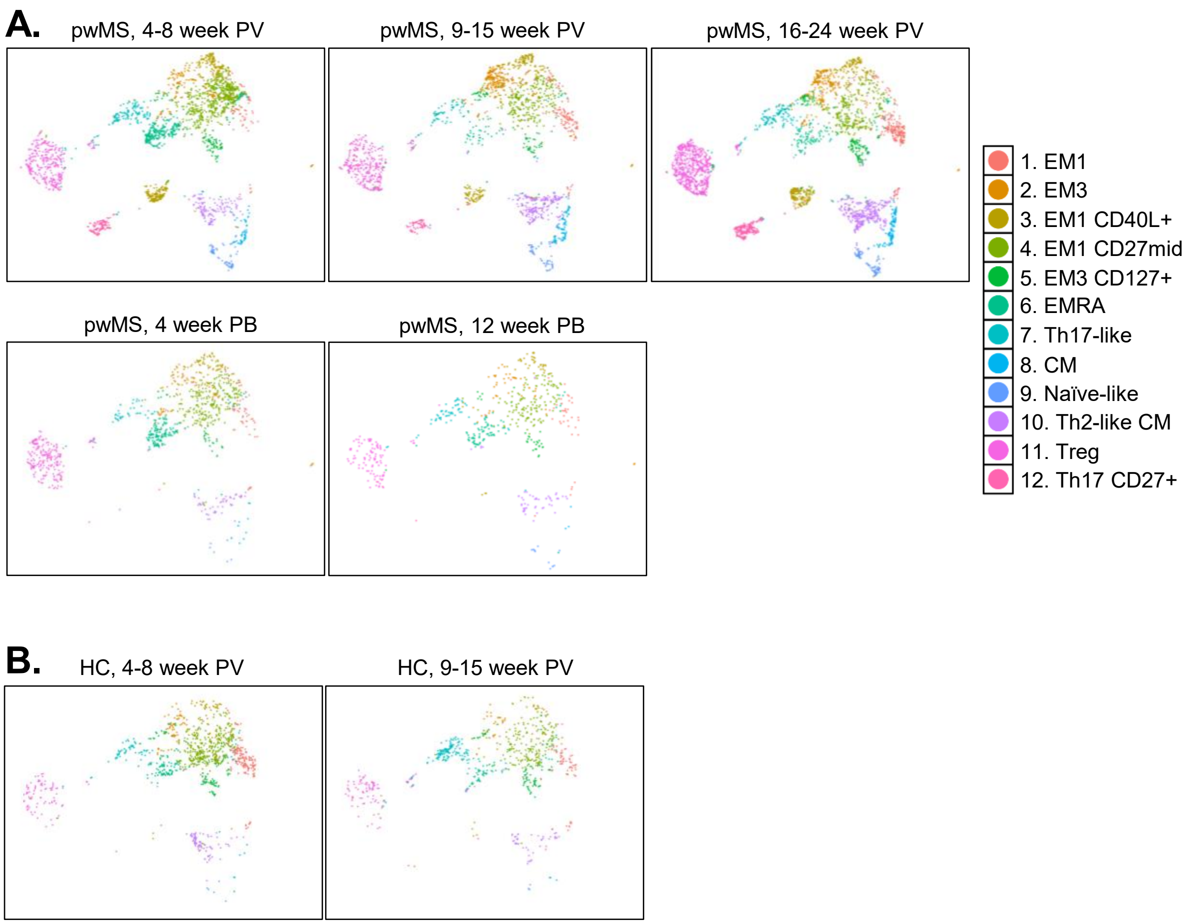


### Supplementary Figure 3

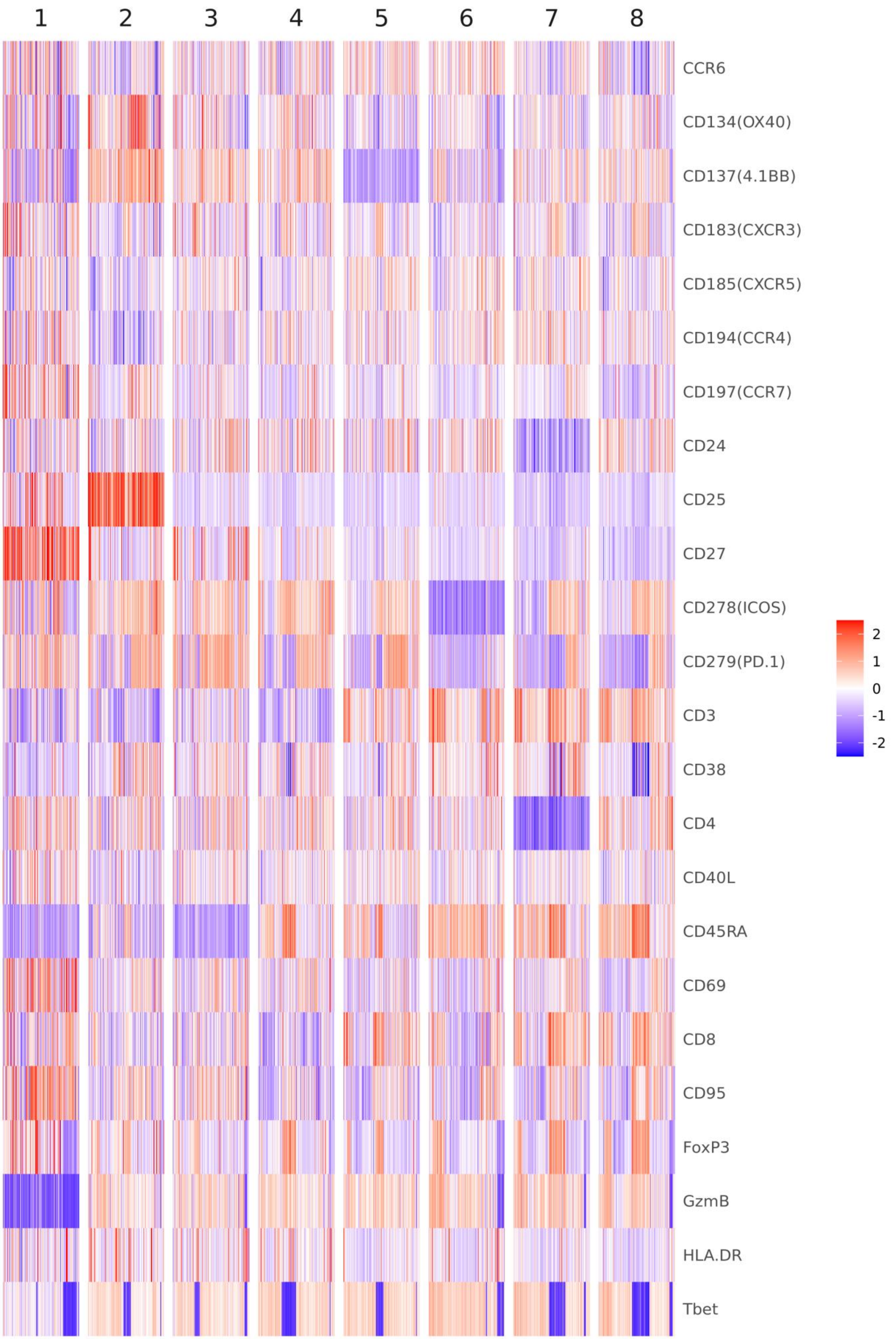


**Supplementary Figure 4**

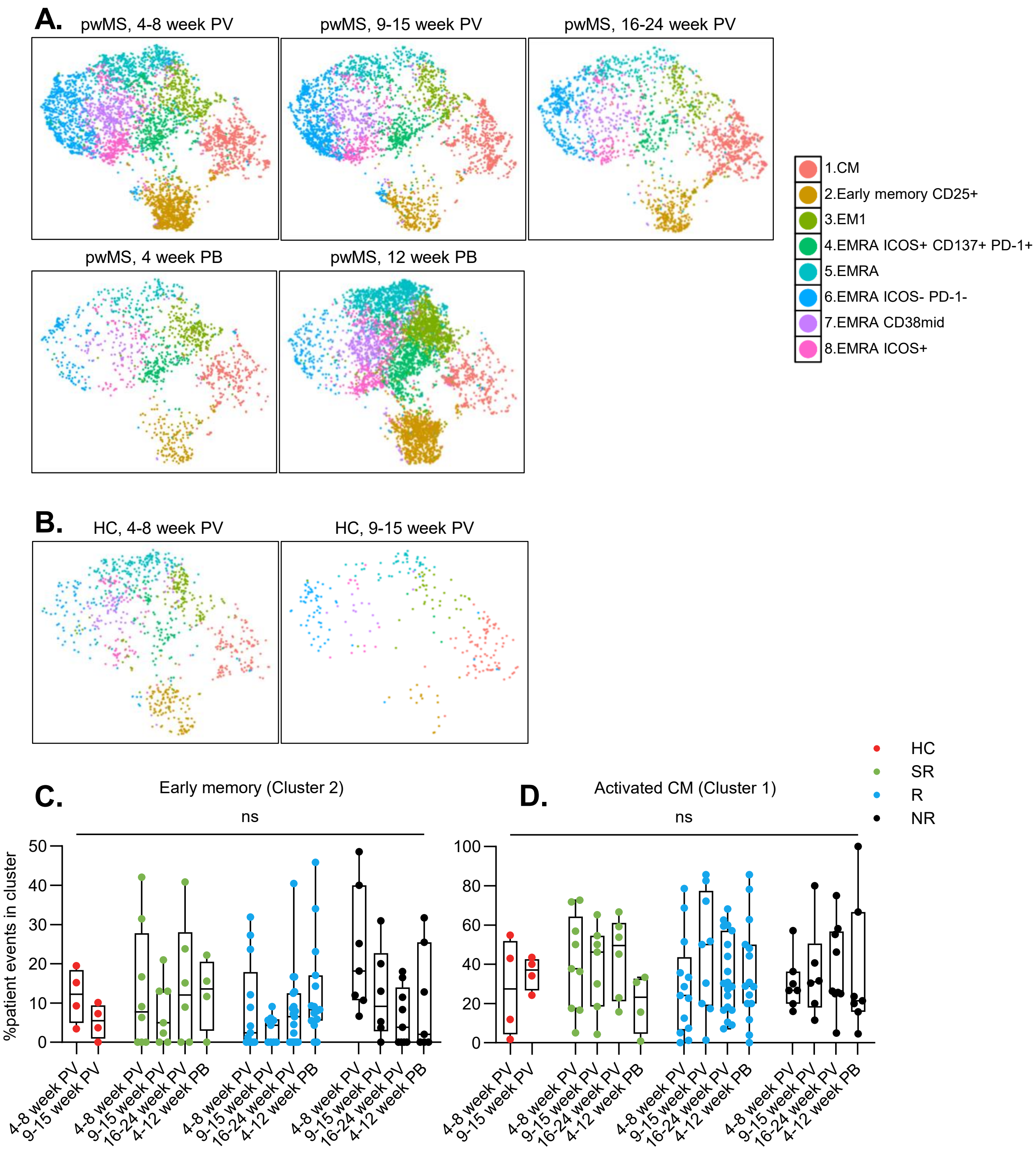


### Supplementary Figure 5

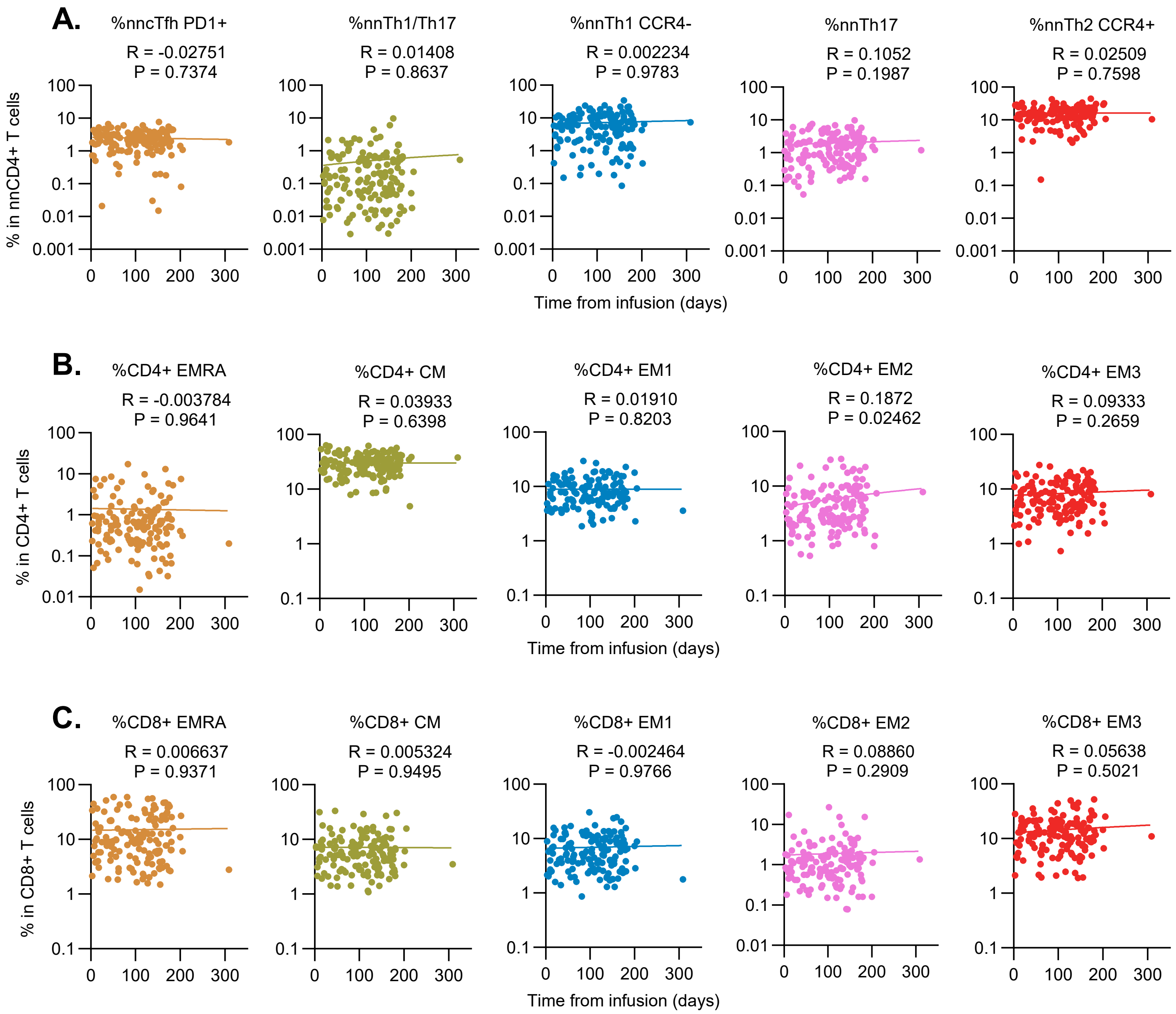


**Supplementary Figure 6**

A.

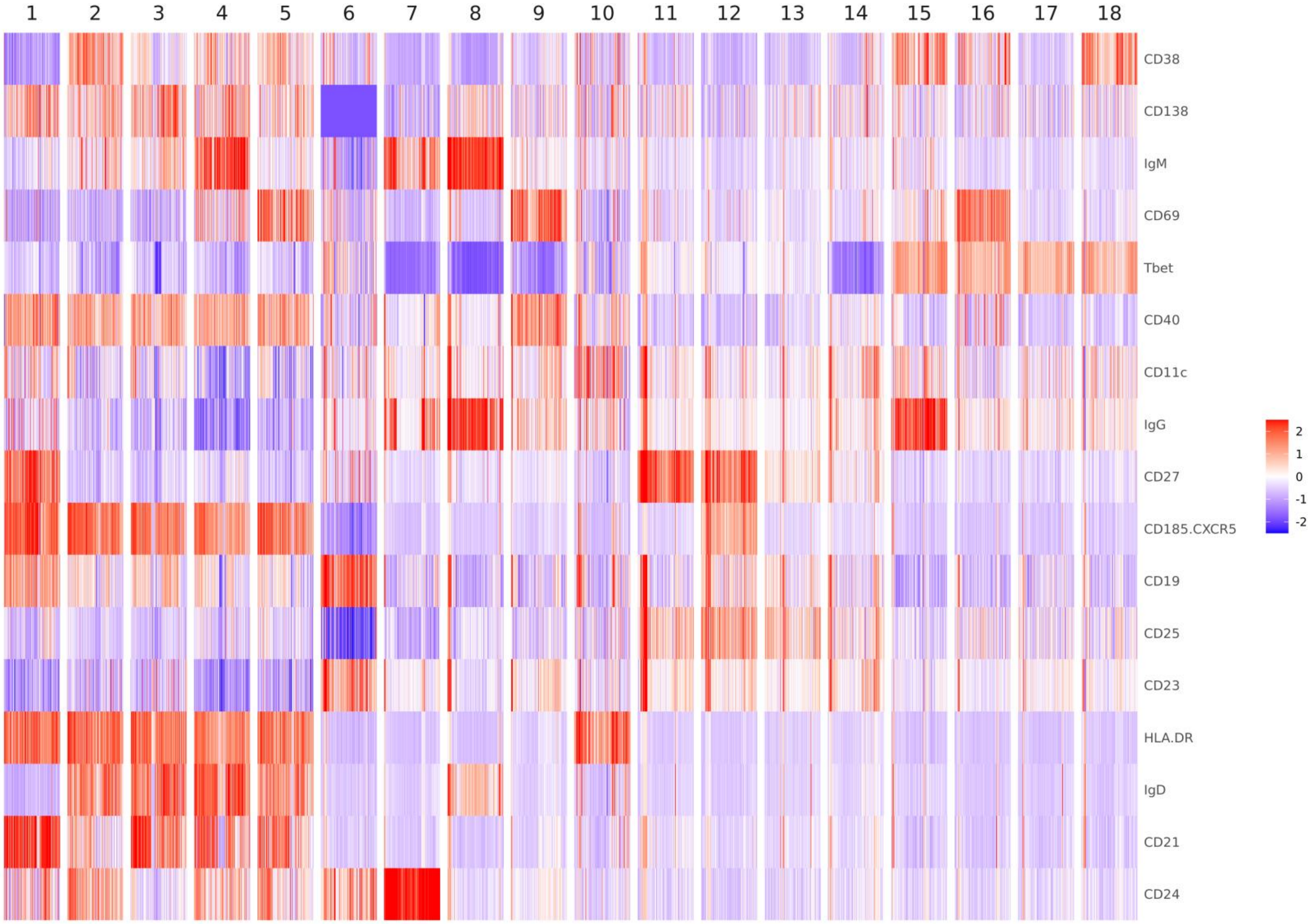


B.

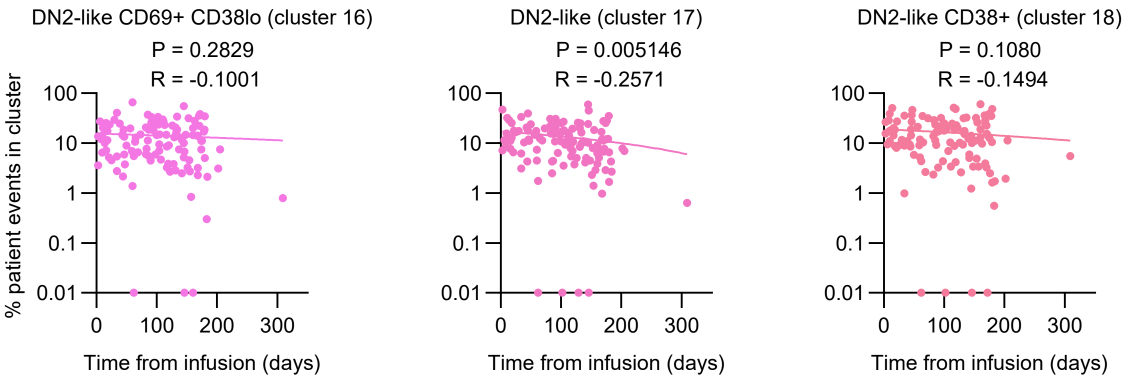


### Supplementary Figure 7
